## Supplementary Information for "Tertiary structure and conformational dynamics of the anti-amyloidogenic chaperone DNAJB6b at atomistic resolution"

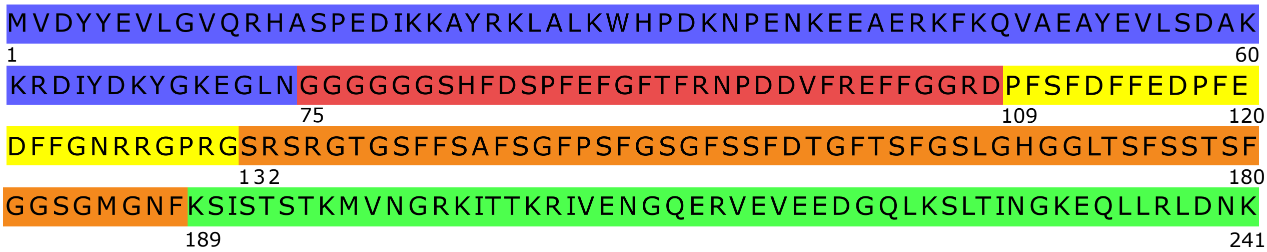


**Supplementary Figure 1: Amino acid sequence of DNAJB6b.** Amino acid sequence of DNAJB6b with each color representing a domain. Blue: J-domain (residue 1 to 74), Red: G/$F_1$ domain (residue 75 to 108), Yellow: G/$F_2$ domain (residue 109 to 131), Orange: S/T domain (residue 132 to 188) and Green: C-terminal domain (CTD) (residue 189 to 241). This color scheme is maintained throughout the text.

| Model | K196 − K29 | K225 − K21 | K225 − K60 | K225 − K70 | Absolute error |
|---|---|---|---|---|---|
| Robetta #1 | 50.47 | 65.95 | 78.05 | 77.73 | 152.20 |
| Robetta #2 | 43.76 | 39.03 | 38.34 | 38.31 | 39.44 |
| Robetta #3 | 9.47 | 20.77 | 40.17 | 38.43 | 40.36 |
| Robetta #4 | 40.16 | 27.99 | 18.75 | 24.34 | 19.07 |
| Robetta #5 | 53.50 | 55.09 | 64.65 | 68.53 | 121.77 |
| RaptorX #1 | 58.57 | 62.86 | 47.64 | 53.41 | 102.48 |
| RaptorX #2 | 72.47 | 57.21 | 40.24 | 43.41 | 93.33 |
| RaptorX #3 | 63.92 | 61.71 | 63.04 | 64.68 | 133.35 |
| RaptorX #4 | 50.94 | 37.88 | 25.48 | 39.85 | 39.19 |
| RaptorX #5 | 61.73 | 53.83 | 48.14 | 59.54 | 103.24 |
| TR-Rosetta #1 | 35.19 | 37.74 | 24.85 | 36.38 | 20.46 |
| TR-Rosetta #2 | 45.25 | 48.20 | 61.10 | 67.99 | 102.54 |
| TR-Rosetta #3 | 53.67 | 43.03 | 46.19 | 55.89 | 78.78 |
| TR-Rosetta #4 | 46.92 | 49.17 | 51.44 | 62.56 | 90.09 |
| TR-Rosetta #5 | 56.01 | 60.26 | 65.88 | 75.79 | 137.94 |
| Alphafold #1 | 24.05 | 40.54 | 54.43 | 56.09 | 63.01 |

**Supplementary Table 1: Homology modelling of DNAJB6b.** Distance between the $C_\alpha$ atoms of the lysines detected in lysine-specific crosslinking mass spectrometry of a DNAJB6 monomer [1]. A distance constraint of 26–30 Å between the $C_\alpha$ atoms of lysines is used to calculate the absolute error [2].

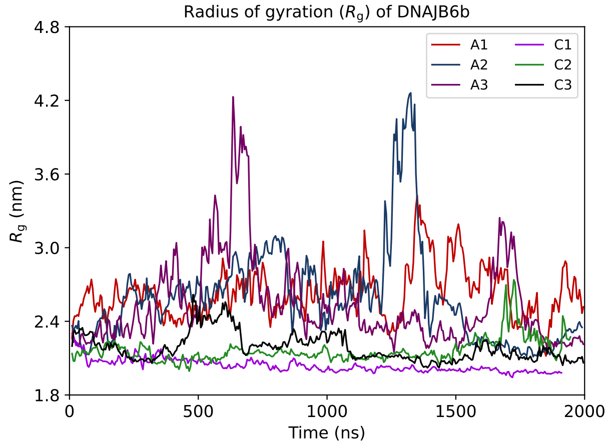


**Supplementary Figure 2: Variation of radius of gyration ($R_g$) with time of DNAJB6b in all the simulations performed.** Variation of the radius of gyration ($R_g$) of DNAJB6b with time. Averaging is performed over 2.5 ns intervals. In simulation A2 and A3, after 1200 ns and 500 ns, respectively, we see a spike in the $R_g$, and this spike indicates the DNAJB6b transitioning from the open state to the extended conformation. A1, A2 and A3 correspond to three replicas using the Amberff99SBdisp force field, whereas C1, C2 and C3 are three replicas performed using the CHARMM36m force field.

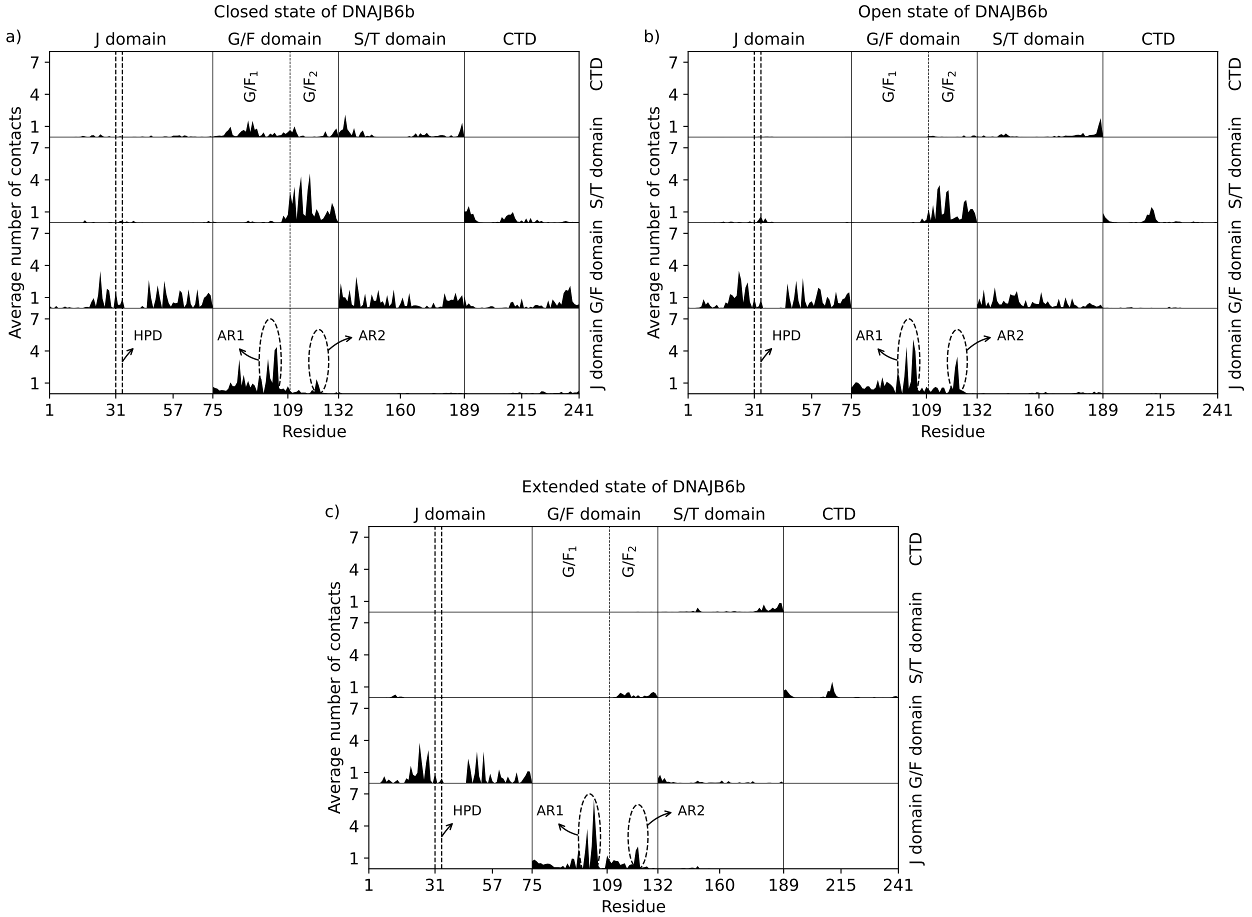


**Supplementary Figure 3: 1D interdomain contact map of DNAJB6b in three different states.** Plots showing the interdomain 1D contact map for the closed **(a)**, open **(b)** and extended **(c)** state of DNAJB6b. The average number of interdomain contacts a residue makes (vertical axis) is plotted against the residue number (horizontal axis) within each interaction domain. The vertical dashed line is drawn at residue 109, separating the G/F domain into the $G/F_1$ and $G/F_2$ domains. The left dashed vertical oval show the interactions between residues 96–104 and the J-domain in anchor region 1 (AR1) and the right dashed oval shows the contacts between residues 118 to 125 and the J-domain in anchor region 2 (AR2). The location of the HPD motif is shown featuring residues 31 to 33.

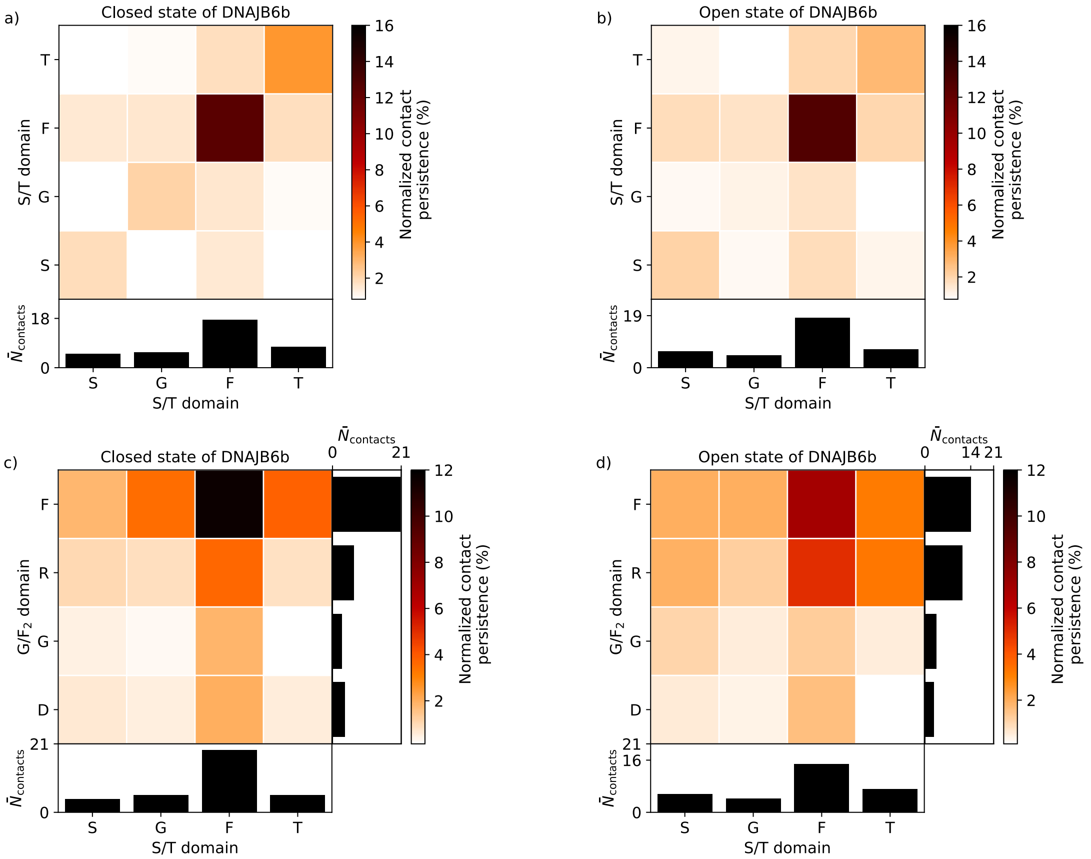


**Supplementary Figure 4: Interactions within the S/T domain and between the S/T and G/F$_2$ domain are predominantly F–F.** Time-averaged number of intramolecular contacts in the S/T domain (termed 'Normalized contact persistence') **(a,b)** and intermolecular contacts between the S/T and G/F$_2$ domain **(c,d)** for the closed and open states as a function of residue type. The contact maps are normalized for residue abundance, where only amino acids that appear at least thrice in the sequence are considered; the bottom panels show the summed contact numbers per residue type ($\bar{N}_{\text{contacts}}$ denote the summations of normalized contacts.). It can be noticed that the predominant inter- and intra- domain interactions are F–F.

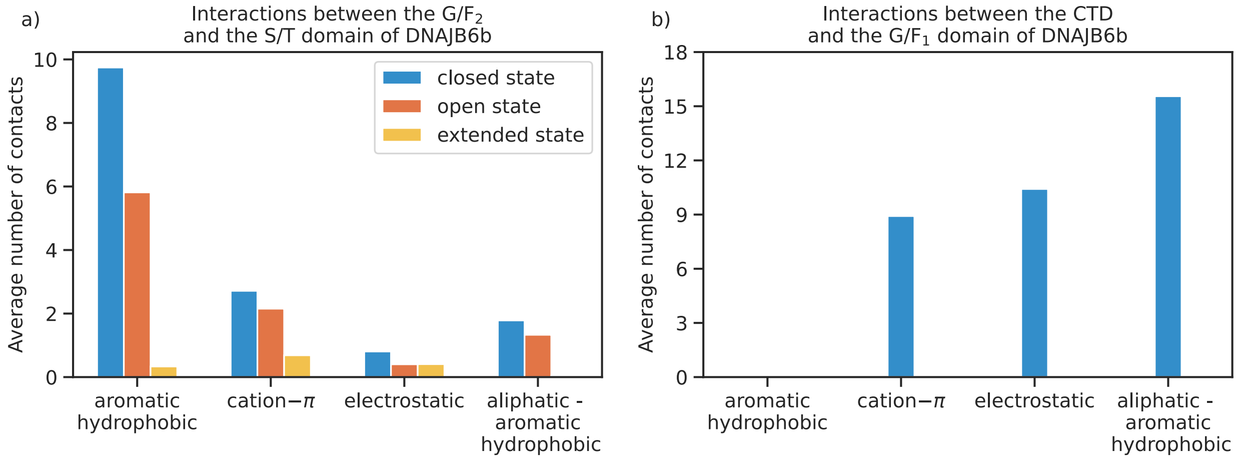


**Supplementary Figure 5: Average number of contacts for different contact types in three different states of DNAJB6b.** Bar plot showing the average number of contacts (averaged over time and averaged over all the simulations performed) for different contact types. **(a)** Contacts between the G/F$_2$ domain (residues 109–131) and the S/T domain (residues 132–189) and **(b)** between the G/F$_1$ domain (residues 75–109) and the CTD (residues 189–241). Aliphatic hydrophobic type contacts are not shown as these are negligible.

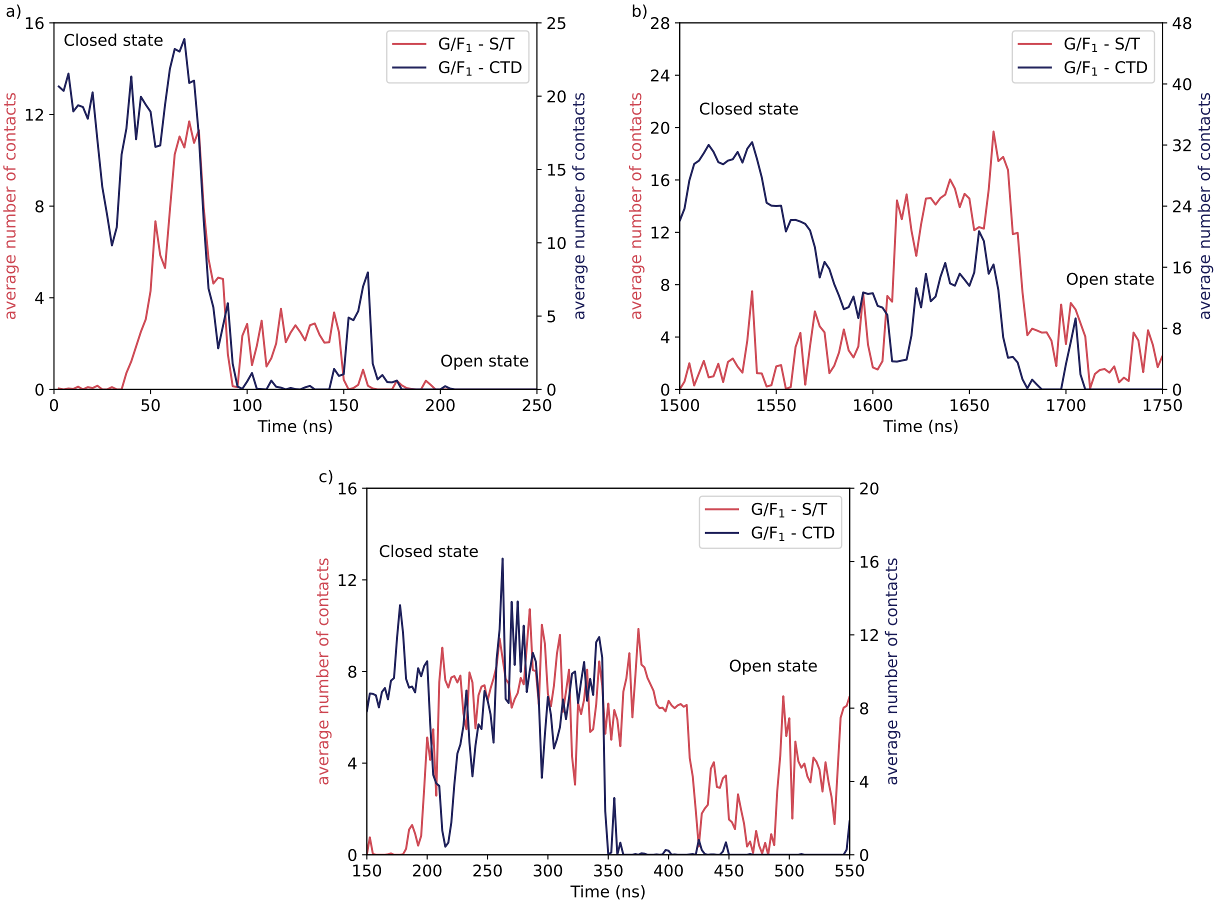


**Supplementary Figure 6: Transition of DNAJB6b from the closed to the open state.** Variation of the average number of contacts varying with time showing the transition of DNAJB6b from the closed state to the open state for the simulations **(a)** C2, **(b)** A2 and **(c)** C3. In simulation C1, we did not observe the transition of DNAJB6b from the closed to the open state and in the simulations A1 and A3, the CTD released from its bound state without the intervention of the S/T domain.

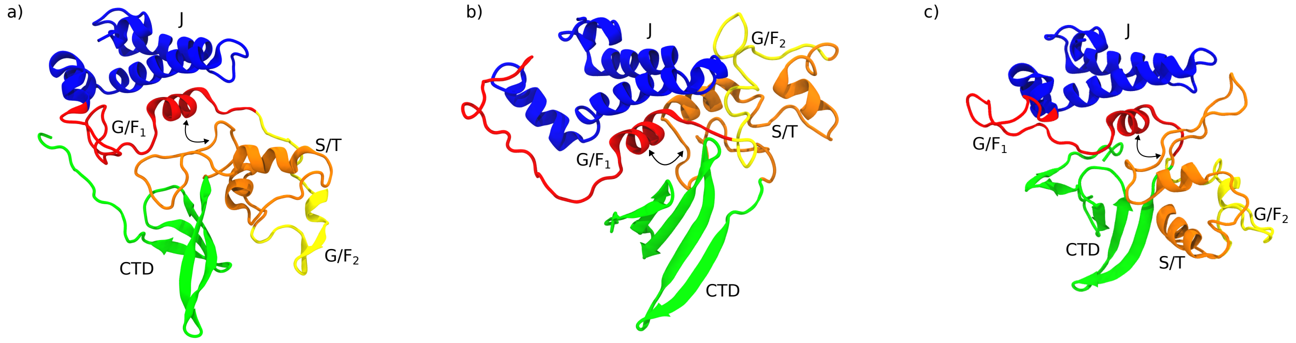


**Supplementary Figure 7: Snapshot showing the interactions between the S/T and G/$F_1$ domain driving transition from the closed to the open state.** Snapshots showing the S/T domain (residues 132 to 188) interacting with the G/$F_1$ domain (residues 75 to 109) (two-head arrow). **(a)** snapshots taken at ∼1650 ns from simulation C2, **(b)** at ∼75 ns from simulation A2 and **(c)** at ∼295 ns from simulation C3. These snapshots show the interdomain interaction between the G/$F_1$ and S/T domain, which triggers DNAJB6b to transition from the closed to the open state. These snapshots can be related to Fig. 6.

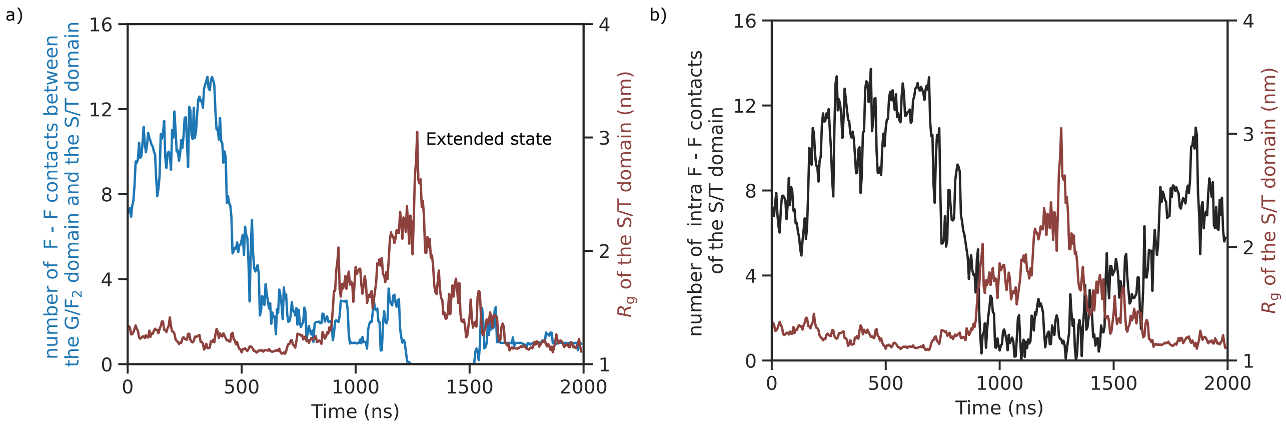


**Supplementary Figure 8: Interactions driving the transition from the open to the extended state.** Plots showing the variation of **(a)** the number of Phe (F)-F interdomain contacts between the G/$F_2$ domain and the S/T domain (left vertical axis), **(b)** the number of F–F intradomain contacts of the S/T domain (left vertical axis) with time. In both (a) and (b) the $R_g$ of the S/T domain is plotted with time to show the effect of inter– and intradomain F–F contacts on the $R_g$ and the transitioning of DNAJB6b from the open to the extended state. In both (a) and (b), $R_g$ and the number of contacts are averaged over 2.5 ns.

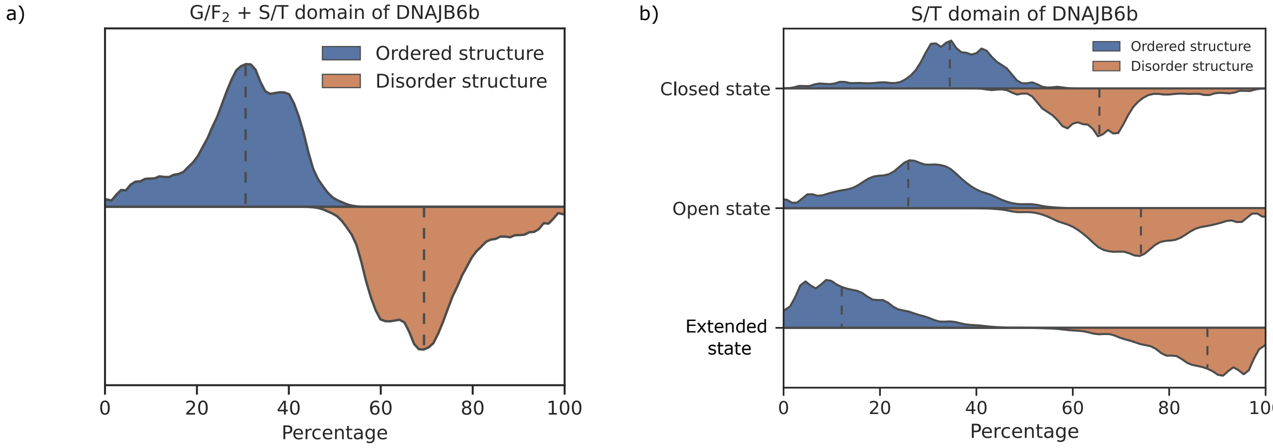


**Supplementary Figure 9: G/$F_2$+S/T domain of DNAJB6b is partly structured.** Horizontal violin plot showing the distribution of ordered and disordered secondary structure percentages of the **(a)** G/$F_2$+S/T domains of DNAJB6b averaged over all the different states and **(b)** S/T domain of DNAJB6b in the closed, open and extended state. The dashed lines in the violin plot represent the median. It can be noticed that the S/T domain is predominantly disordered in the extended state of DNAJB6b, which is not the case in the open and closed state of DNAJB6b. Helix, β-sheet, $3_{10}$-helix, pi-helix and isolated bridge are considered to be ordered, turn and coil are considered to be disordered for calculating the distributions. The median percentages of the disordered secondary structure of the S/T domain in the closed, open and extended states are 66, 74 and 86 %, respectively.

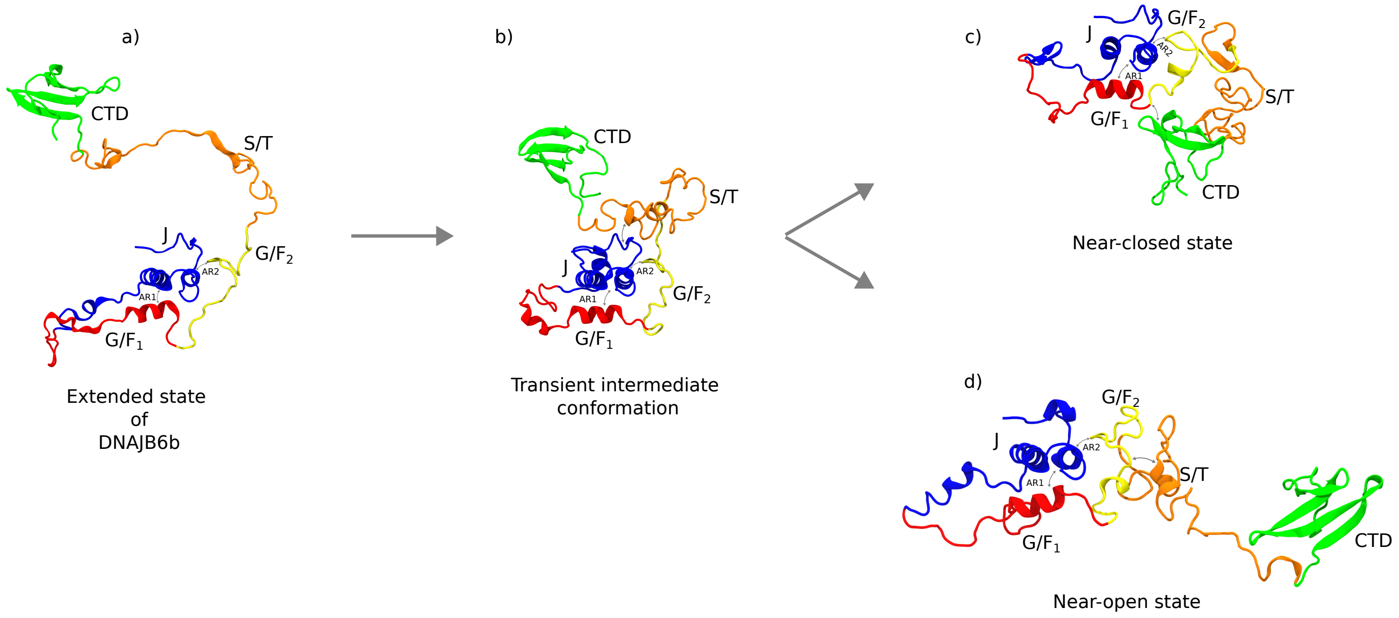


**Supplementary Figure 10: DNAJB6b changing its conformation from the extended state to the near closed or open state.** Snapshots showing the transitioning of the **(a)** extended state of DNAJB6b on its way towards the closed and open state through a **(b)** transient intermediate conformation in which the S/T domain interacts with helix I of the J-domain. This transient state is further converted to a **(c)** near-closed state in which the CTD does not yet interact with the $G/F_1$ domain (but instead interacts with the N-terminal of the $G/F_2$ domain) or to the **(d)** near-open state in which the S/T domain is not yet fully collapsed as in the open state .

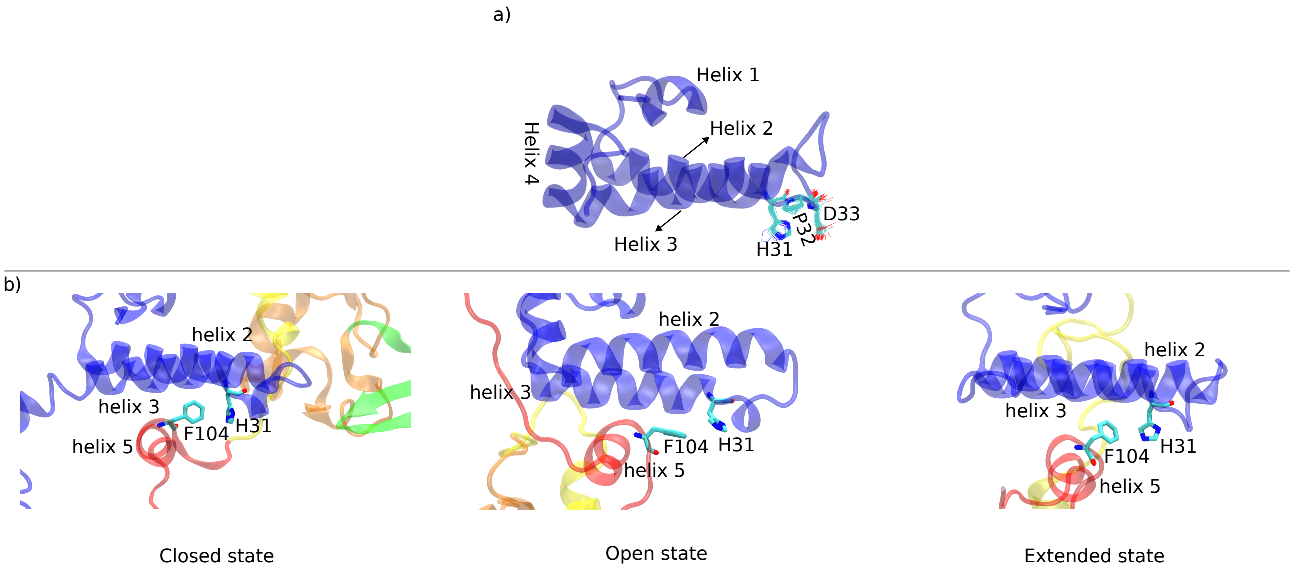


**Supplementary Figure 11: Location of the HPD motif and its interaction with F104 of helix V. (a)** Snapshot showing the J-domain (in secondary structure representation) and the averaged location of the HPD motif (shown in line representation), and **(b)** snapshots showing the prominent interactions observed between H31 of the HPD motif and F104 of helix V for the closed, open and extended states. Residue F104 interacts with histidine 31 (H31) via an NH···π hydrogen bond of the imidazole benzene ring [3–8].

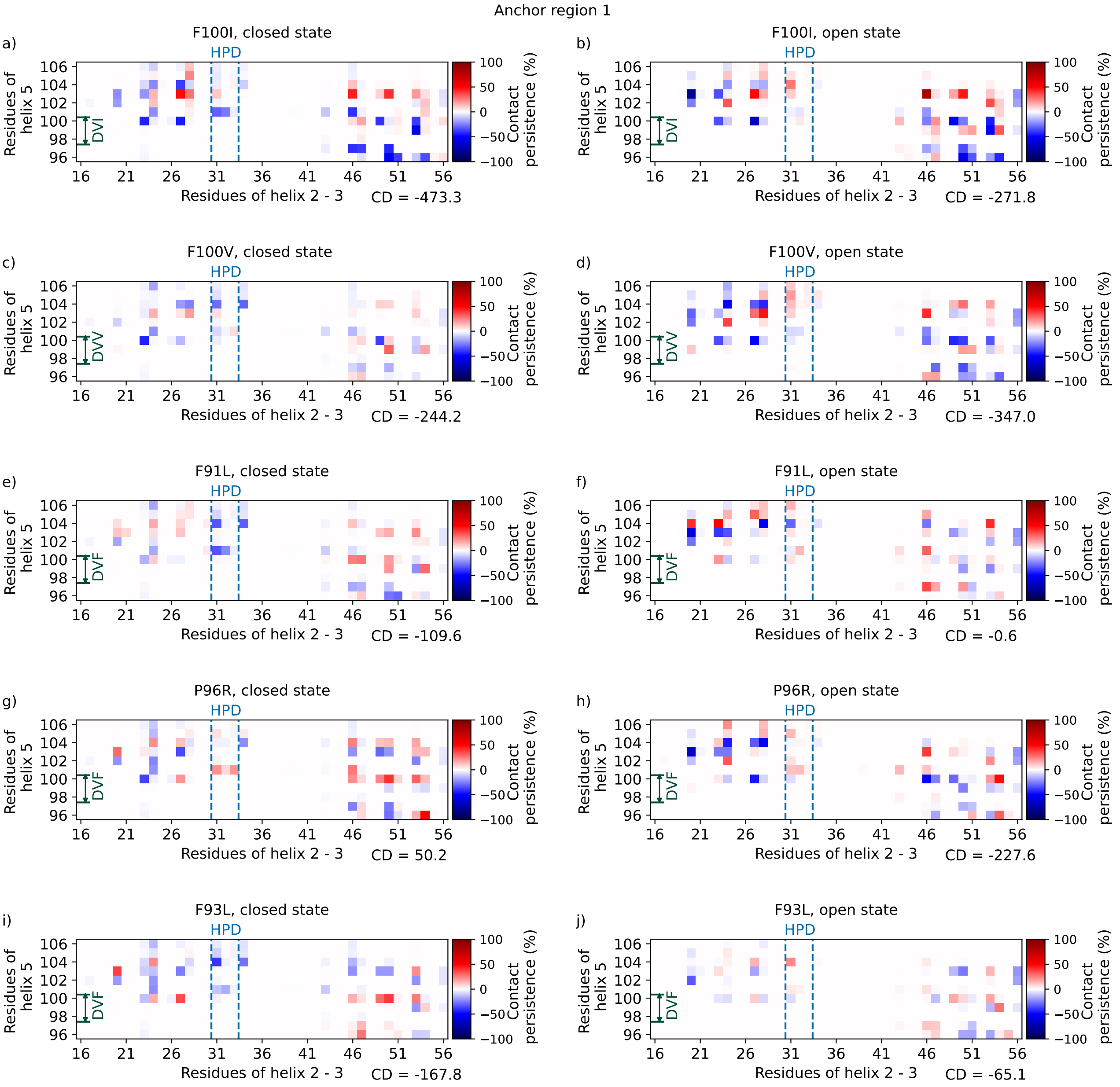


**Supplementary Figure 12: LGMD mutations cause a decrease in contacts between helix V and helices II/III.** Contact map of anchor region 1 (AR1) showing the increase or decrease in contacts for the LGMDD1 mutated DNAJB6b compared to wild-type DNAJB6b. Contact maps are computed for F100I, F100V, F91L, P96R and F93L in the closed state **(a, c, e, g and i)** and open state **(b, d, f, h and j)**. Positive values indicate increased, and negative values indicate decreased interactions within anchor region 1. A cumulative difference (CD, %) between mutated and WT DNAJB6b in contact persistence for anchor region 1 is reported for every contact map. The CD is determined by summing the contact persistence values for all observed interresidue interactions in the contact map. In the scenario where the wild type DNAJB6b has no contacts but the LGMDD1 mutated DNAJB6b does, the percentage will solely reflect the mutated DNAJB6b, denoted with a positive sign to indicate the presence of contacts compared to the wild type. Conversely, if the wild type DNAJB6b has contacts that are absent in the LGMDD1 mutated DNAJB6b, the percentage will solely reflect the wild type DNAJB6b, denoted with a negative sign to indicate the absence of contacts compared to the wild type. It can be observed that the CD is negative for all the LGMD-associated mutations except for P96R (closed state), indicating that almost all mutations on the G/F domain will lead to a decrease in contacts between helix V and helices II/III. Despite the complete destabilization of AR1 observed in the P96R open state, it is not evident in the contact map since the map is constructed over the entire simulation duration.

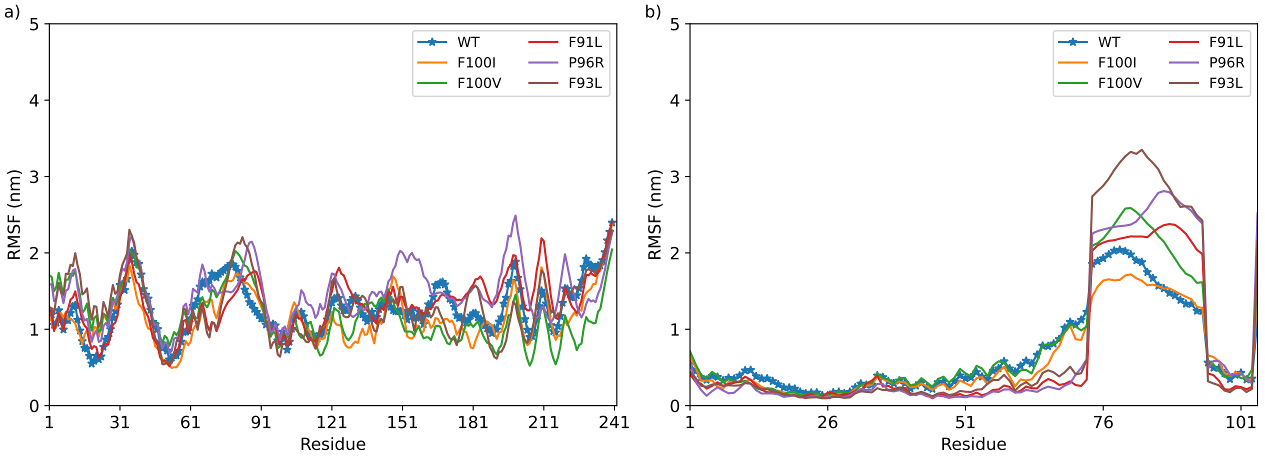


**Supplementary Figure 13: Effect of LGMDD1 mutations on the residue fluctuations in DNAJB6b.** Root mean square fluctuations (RMSF) are plotted for the open state of DNAJB6b and the LGMDD1-associated point mutations. The RMSF are calculated with the alignment on **a)** the entire protein's backbone, **b)** the J-domain (resid 1 – 74) and the helix V (resid 96–104).

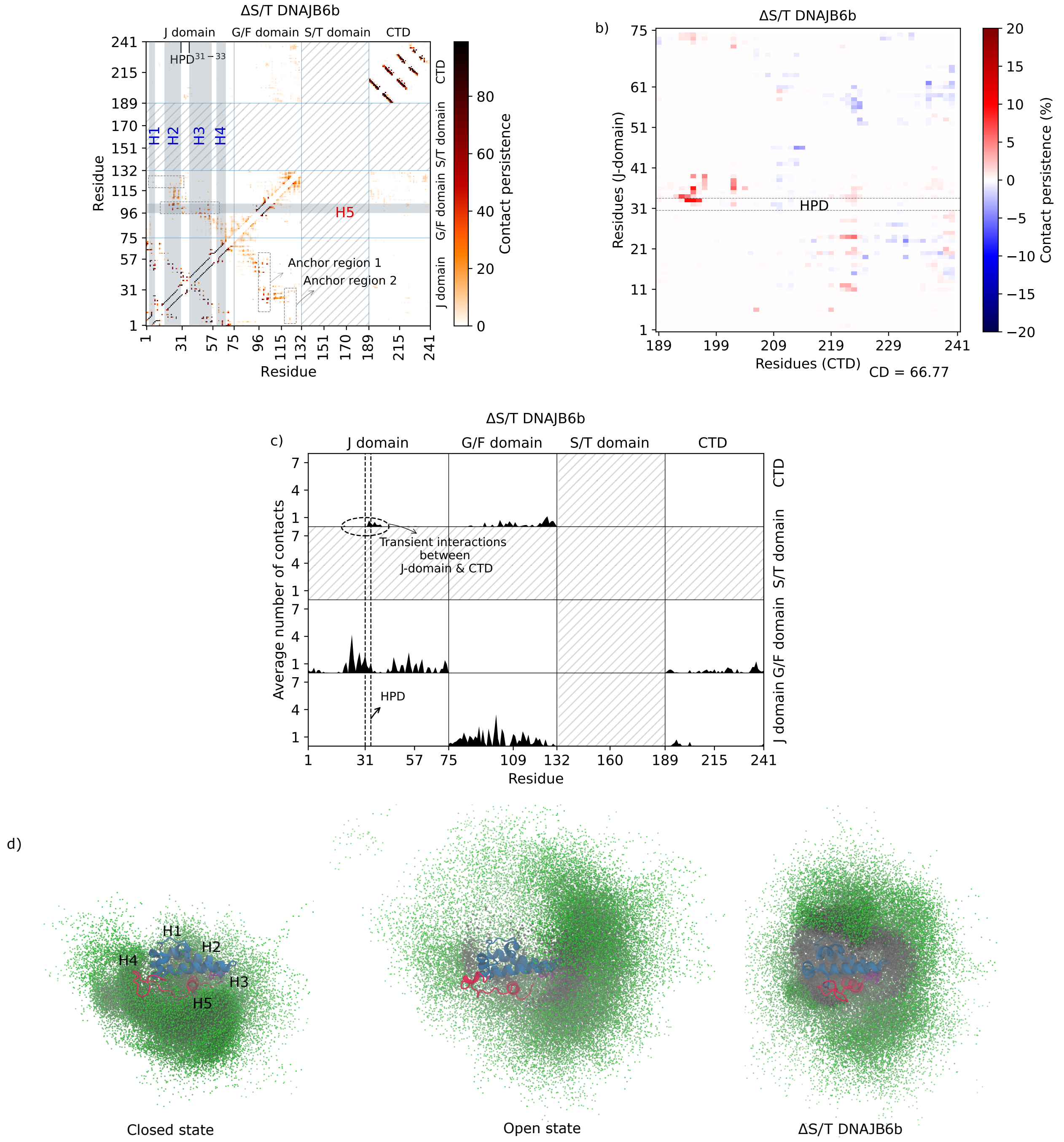


**Supplementary Figure 14: Interdomain interactions for the ΔS/T DNAJB6b construct is different compared to full-length DNAJB6b. (a)** Contact map of ΔS/T DNAJB6b. The color bar corresponds to the percentage of time the contact existed during the simulation. The contact map is divided into 4 sections according to the domains in DNAJB6b. The J-domain contains 4 helices shown by vertical gray bars. The horizontal gray bar shows the helix V of the $G/F_1$ domain. The contacts enclosed in the dashed rectangle indicate the anchor regions 1 and 2 (AR1 and AR2). It can be noticed that there are no contacts in anchor region 2, **(b)** Contact map between the CTD and J-domain showing the increase or decrease in contacts for ΔS/T DNAJB6b compared to wild-type DNAJB6b. Positive values indicate increased and negative indicate decreased interactions. The location of the HPD motif is shown featuring residues 31 to 33. A cumulative difference (CD, %) between ΔS/T and WT DNAJB6b in contact persistence is reported. The CD is determined by summing the contact persistence values for all observed interresidue interactions in the contact map. It can be noticed that there is a 5 % increase in contacts between the CTD and the N-terminal of the J-domain compared to wildtype, while there is a 10–15 % increase in interactions between the CTD and the HPD motif for Δ S/T DNAJB6b compared to full-length DNAJB6b, **(c)** Plot showing the interdomain 1D contact map for ΔS/T DNAJB6b. The average number of interdomain contacts a residue makes (vertical axis) is plotted against the residue number (horizontal axis) within each interaction domain. The location of the HPD motif is shown featuring residue 31 to 33. The horizontal oval shows the transient interdomain interactions between the J-domain and the CTD, (caption continued on next page)

(Supplementary Figure 14, caption continued) **(d)** Snapshots showing the conformational space sampled by the C-alpha atoms of the CTD relative to the J domain for the closed and open states of full-length DNAJB6b and for ΔS/T DNAJB6b. For comparison, 1000 snapshots, equally spaced in time, were chosen to show the conformational space sampled by the CTD. The distribution of the CTD within 10 Å of the J-domain were shown in black. It can be noticed from these snapshots that, in case of ΔS/T DNAJB6b, the CTD has many interactions with the J-domain compared to full-length DNAJB6b. Alignment is done on the J-domain prior to plotting the distribution of $C_\alpha$ atoms of the CTD.
